# BiteOscope: an open platform to study mosquito blood-feeding behavior

**DOI:** 10.1101/2020.02.19.955641

**Authors:** Felix JH Hol, Louis Lambrechts, Manu Prakash

**Affiliations:** Department of Bioengineering, Stanford University, Stanford, CA, USA; Insect-Virus Interactions Unit, Institut Pasteur, UMR2000, CNRS, Paris, France; Center for research and Interdisciplinarity, Université de Paris, Paris, France

**Author notes:** For correspondence (FJHH).

## Abstract

Female mosquitoes need a blood meal to reproduce, and in obtaining this essential nutrient they transmit deadly pathogens. Although crucial for the spread of mosquito-borne diseases, our understanding of skin exploration, probing, and engorgement, is limited due to a lack of quantitative tools. Indeed, studies often expose human subjects to assess biting behavior. Here, we present the biteOscope, a device that attracts mosquitoes to a host mimic which they bite to obtain an artificial blood meal. The host mimic is transparent, allowing high-resolution imaging of the feeding mosquito. Using machine learning we extract detailed behavioral statistics describing the locomotion, pose, biting, and feeding dynamics of *Aedes aegypti, Aedes albopictus, Anopheles stephensi*, and *Anopheles coluzzii*. In addition to characterizing behavioral patterns, we discover that the common insect repellent DEET repels *Anopheles coluzzii* upon contact with their legs. The biteOscope provides a new perspective on mosquito blood feeding, enabling high-throughput quantitative characterization of the effects physiological and environmental factors have on this lethal behavior.

## Introduction

Blood feeding is essential for mosquito reproduction, and in the process mosquitoes transmit myriad pathogens to their (human) host. Yet despite being the focal point of pathogen transmission, many aspects of blood feeding remain ill understood. The initial step in obtaining a blood meal, flying towards a host, is relatively well characterized (***Dekker and Cardé, 2011***; ***McMeniman et al., 2014***; ***van Breugel et al., 2015***). The steps that unfold after a mosquito has landed on a host, however, are much less understood. Once landed, mosquitoes exhibit exploratory bouts during which the legs and proboscis frequently contact the skin (***Jones and Pilitt, 1973***; ***De Jong and Knols, 1995***; ***Clements, 2013***). An increasing body of literature reports the presence of receptors involved in contact-dependent sensing on the legs and proboscis (***Sparks et al., 2013***; ***Matthews et al., 2019***; ***Dennis et al., 2019***), suggesting that these appendages evaluate the skin surface and thus serve an important role in bite-site selection. Yet the role and mechanism of contact-dependent sensing in blood feeding, is largely unclear (***Benton, 2017***). In addition to the body parts that come in contact with the skin surface, the skin piercing labrum also serves as a chemosensory organ, guiding blood feeding in currently unknown ways (***Lee, 1974***; ***Werner-Reiss et al., 1999***; ***Jové et al., 2020***).

In addition to external cues, an animal’s (internal) physiology may also affect its behavior. Nutrition, hydration, and pathogen infections, for instance, have been hypothesized to affect blood feeding behavior, e.g. by altering feeding avidity (i.e. number of feeding attempts) or the size of the meal taken (***Rossignol et al., 1984***; ***Choumet et al., 2012***; ***Cator et al., 2013***; ***Vantaux et al., 2015***; ***Hagan et al., 2018***). These topics, however, remain a matter of debate, due to a lack of (standardized) assays to measure mosquito behavior (***Stanczyk et al., 2017***). Quantitative mapping of *Drosophila* behavior provides an important perspective, suggesting that innovative experimental approaches and computational tools can fuel the acquisition of new insights (e.g. ***Itskov et al. (2014***); ***Branson et al. (2009***); ***Kain et al. (2013***); ***Moreira et al. (2019***); ***Berman et al. (2014***); ***Kabra et al. (2013***)). Yet apart from olfactometers and other flight chambers, very few assays to characterize the blood-feeding behavior of mosquitoes exist (***Geier and Boeckh, 1999***; ***Verhulst et al., 2011***; ***McMeniman et al., 2014***; ***van Breugel et al., 2015***). Due to this paucity of assays, studies often expose human subjects to quantify the number of landings and/or bites, or the time it takes to complete a blood meal, and score experimental outcomes by hand (***Jones and Pilitt, 1973***; ***Ribeiro, 2000***; ***Moreira et al., 2009***; ***DeGennaro et al., 2013***; ***Dennis et al., 2019***). The use of humans as bait constrains the number and type of experiments that can be done (e.g. prohibiting the use of infected mosquitoes) and limits the type, detail, and throughput of measurements that can be made. Furthermore, the opaque nature of skin prevents the visualization of the stylets after piercing the skin leaving this aspect of blood feeding almost entirely unstudied, except for one notable study using intravital imaging of dissected mouse skin (***Choumet et al., 2012***) and two much earlier descriptions (***Gordon and Lumsden, 1939***; ***Griffiths et al., 1952***).

To overcome these limitations, we developed the biteOscope, an open platform that allows the high-resolution and high-throughput characterization of surface exploration, probing, and engorgement by blood feeding mosquitoes. The biteOscope consists of a rudimentary skin mimic: a substrate that attracts mosquitoes to its surface, induces them to land, pierce the surface, and engage in blood feeding. The bite substrate can be mounted in the wall of a mosquito cage allowing freely behaving mosquitoes access. By virtue of its transparent nature, the substrate facilitates imaging of mosquitoes interacting with it, including the visualization of the skin piercing mouthparts of the mosquito. We developed a suite of computational tools that automates the extraction of behavioral statistics from image sequences, and use machine learning to track the individual body parts of behaving mosquitoes. These capabilities enable a detailed characterization of blood feeding mosquitoes. We demonstrate that the biteOscope is an effective instrument to study the behavior of several medically relevant species of mosquito and describe behavioral patterns of the two main vectors of dengue, Zika, and chikungunya virus (*Aedes aegypti* and *Aedes albopictus*), and two important malaria vectors (*Anopheles coluzzii* and *Anopheles stephensi*). The biteOscope allows detailed tracking of the complex interactions of mosquitoes with a substrate, and can be used to characterize behavioral alterations in the presence of chemical surface patterns. Using this capability, we provide evidence that DEET repels *Anopheles coluzzii* upon contact with their legs, demonstrating the utility of body part tracking to understand behaviors mediated by contact-dependent sensing. We anticipate that the biteOscope will enable studies that increase our understanding of the sensory biology and genetics of blood feeding, and the effects external (environmental) and internal (physiology) variables have on this behavior. Given its relevance for pathogen transmission, dissecting the interplay between the mosquito sensory system and host-associated cues during blood feeding is of clear interest, and may suggest new avenues to interfere with blood feeding, and eventually curb pathogen transmission.

## Results

### The biteOscope

To allow mosquitoes to engage in blood feeding and feed to full repletion, a device needs to attract mosquitoes, allow them to explore and pierce the surface, and subsequently imbibe a blood meal. To design a tool that can easily be used in a variety of ‘mosquito labs’ (including (semi-)field settings), we sought to recapitulate this behavioral sequence using readily available and low-cost laboratory materials. Heat is a dominant factor in short-range mosquito attraction and can be used to attract mosquitoes to a surface and elicit probing behavior (***Healy et al., 2002***; ***Corfas and Vosshall, 2015***; ***Zermoglio et al., 2017***). We constructed a bite substrate using an optically clear flask filled with water as a controllable heat source (see Fig. 1A). An artificial blood meal is applied on the outside of the flask and covered using Parafilm (a commonly used membrane in laboratory blood feeders) creating a thin fluid cell on which mosquitoes can feed. To elicit blood feeding in a transparent medium, we use adenosine triphosphate (ATP) as a strong phagostimulant, which, together with an osmotic pressure similar to that of blood and the presence of sodium ions, is su*ffi*cient to induce mosquitoes to feed to full engorgement (***Galun et al., 1963***; ***Duvall et al., 2019***).

**Figure 1.**
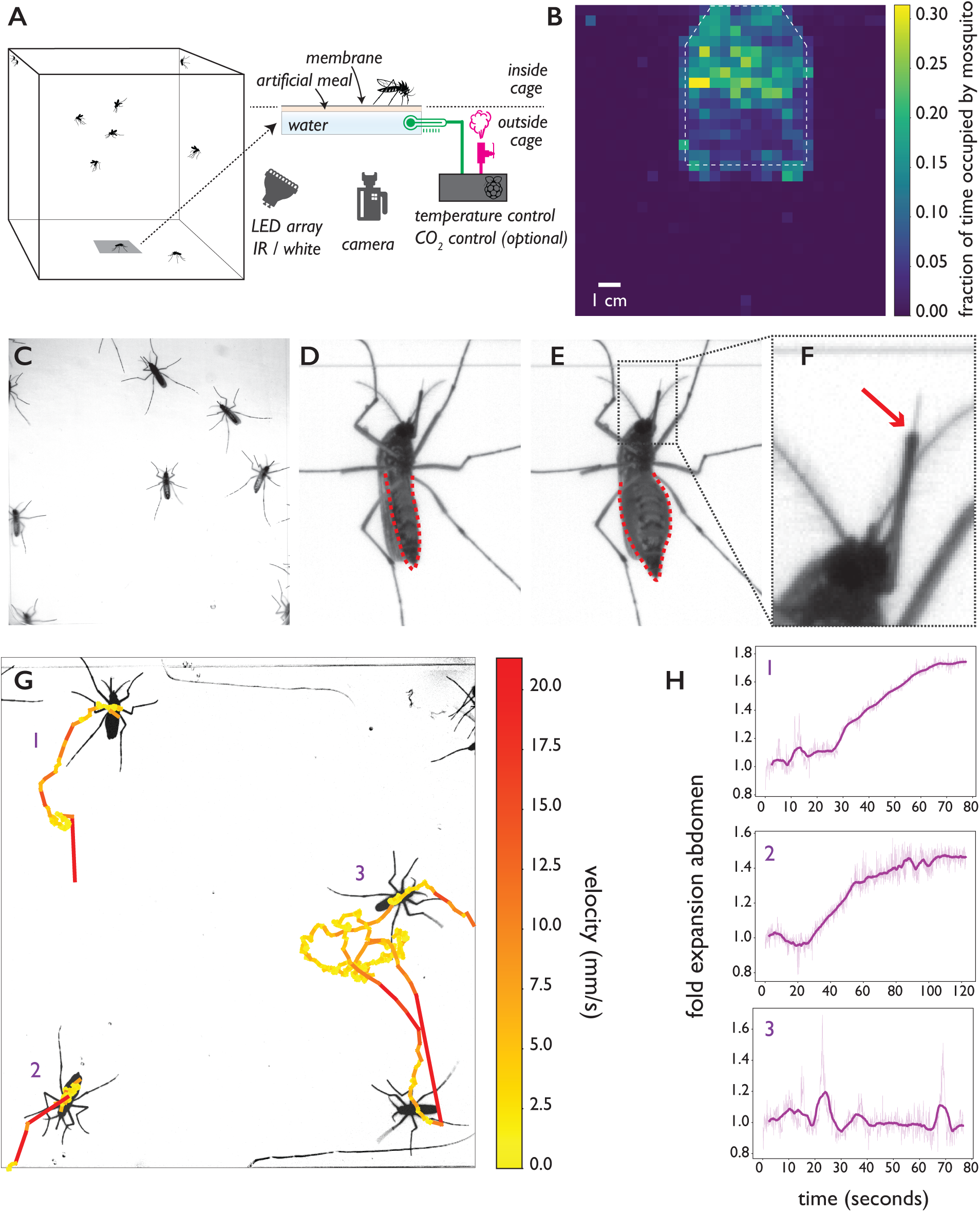
The biteOscope. (**A**) Schematic of the set up. The bite substrate consists of a water bath (cell culture flask) that is mounted in the floor or wall of a cage, allowing freely flying mosquitoes access. An artificial meal is applied on the outside surface of the culture flask and covered using a Parafilm membrane, water in the flask is temperature controlled using a Raspberry Pi reading a temperature probe, and a Peltier element for heating (0.1 °C accuracy). The Raspberry Pi optionally controls the inflow of gas. Illumination is provided by an array of white or IR LED. A camera and lens situated outside the cage images mosquitoes (abdominal view) through the bite substrate. (**B**) Two-dimensional histogram (heatmap) showing mosquito presence on the bite substrate (indicated with a dashed line) and on the surrounding wall. Mosquitoes spend more time on the bite surface. (**C**) Raw image of *Ae. aegypti* on the bite substrate. (**D - F**) Images of an *Ae. aegypti* mosquito that has pierced the membrane and inserted its stylet into the meal. After imbibing, the abdomen dilates. The red arrow in (**F**)) indicates the tip of the labium where the stylets (visible as a thin needle-like structure) pierce the surface and enter the artificial meal. (**G**) Tracks showing movement of *Ae. aegypti* on the bite substrate, color of tracks indicates velocity. (**H**) Fold expansion of the abdomen over time, indicating full engorgement in mosquitoes 1 and 2, and no feeding in mosquito 3 of panel (G).

To allow freely behaving mosquitoes access to the bite substrate, we constructed acrylic cages having an opening in the wall or floor where the bite substrate can be mounted. The bite substrate is transparent, facilitating imaging with a camera mounted outside the cage (Fig. 1A shows a schematic of the set up). For the majority of data presented here, we used a 4.3 × 4.3 cm field of view (see Fig. 1C) which allows up to 15 mosquitoes to explore and feed simultaneously while providing images at a resolution where small body parts like the stylets can easily be resolved. Depending on experimental requirements, the field of view (and correspondingly assay throughput) can be much larger at the expense of resolution. Figure 1B, for example, shows a 13 × 13 cm field of view. Individual mosquitoes can be easily tracked at that resolution, yet the visualization of small body parts is challenging. Experiments on *Ae. aegypti* and *Ae. albopictus*, both active during the day, were performed using white light illumination; we used an infrared (IR) LED array as light source during experiments on *An. coluzzii* and *An. stephensi* which were performed in the dark, corresponding to their peak activity during the night. Figure 1B demonstrates that *Ae. aegypti* mosquitoes show strong attraction to the bite substrate (surface indicated using a dashed line) and spend more time on its surface compared to the surrounding wall. Figure 1C-F shows *Ae. aegypti* undertaking the full blood feeding trajectory on the substrate: starting with surface exploration (Fig. 1C and G), piercing of the membrane and insertion of the stylet into the artificial meal (Fig 1D-F), and feeding to full engorgement, as evidenced by the expanded abdomen (Fig. 1E). Videos 1, 2, 3, and 4 show blood feeding *Ae. albopictus, Ae. aegypti, An. stephensi*, and *An. coluzzii*, respectively. Imaging the stylet at high temporal and spatial resolution (videos 1 and 5) as it evaluates the artificial meal reveals the striking dexterity of the organ as it rapidly bends, extends, and retracts—aspects of feeding that normally remain hidden inside the skin.

### Automatic characterization of the blood feeding behavior of multiple species

We created a computational pipeline to extract behavioral statistics from image sequences (see Fig. S1 for an overview and Materials & Methods for details). The position of individual mosquitoes is tracked over time to yield locomotion statistics (see Fig. 1G and Video 6), and select all time slices that make up a single behavioral trajectory (e.g. landing, exploration, feeding, and take off). To determine a mosquito’s engorgement status, we take advantage of the dilation of the mosquito abdomen when it takes a blood meal (Fig. 1E). We determine a mosquito’s body shape (excluding appendages) using an active contour model to quantify feeding dynamics and engorgement status at each timepoint of a trajectory, and define full engorgement as the saturation of abdominal dilation (see Fig. 1G1-3 and Video 7). Together with locomotion statistics, engorgement data provides a high-level description of the behavioral trajectory.

To assess the capability of the biteOscope to characterize the behavior of different species of mosquito, we performed experiments with the two most important vectors of arboviral diseases (*Ae. aegypti* and *Ae. albopictus*) and two dominant malaria vectors (*An. stephensi* and *An. coluzzii*, formerly known as *Anopheles gambiae* M molecular form). Figures 2 and S2 show locomotion and feeding statistics for the four species. All species land readily on the bite substrate and undertake exploratory bouts leading to full engorgement in 18%, 7%, 4%, and 14% of all trajectories and 46%, 22%, 10%, and 31% of all > 10 second trajectories, for *Ae. aegypti, Ae. albopictus, An. stephensi*, and *An. coluzzii*, respectively, when offered a meal consisting of 1 mM ATP in phosphate buffered saline (PBS). Figure 2A-D shows summary statistics of 350 behavioral trajectories of *An. coluzzii* obtained from 1 hour and 15 minutes of imaging data, demonstrating the throughput of the biteOscope.

**Figure 2.**
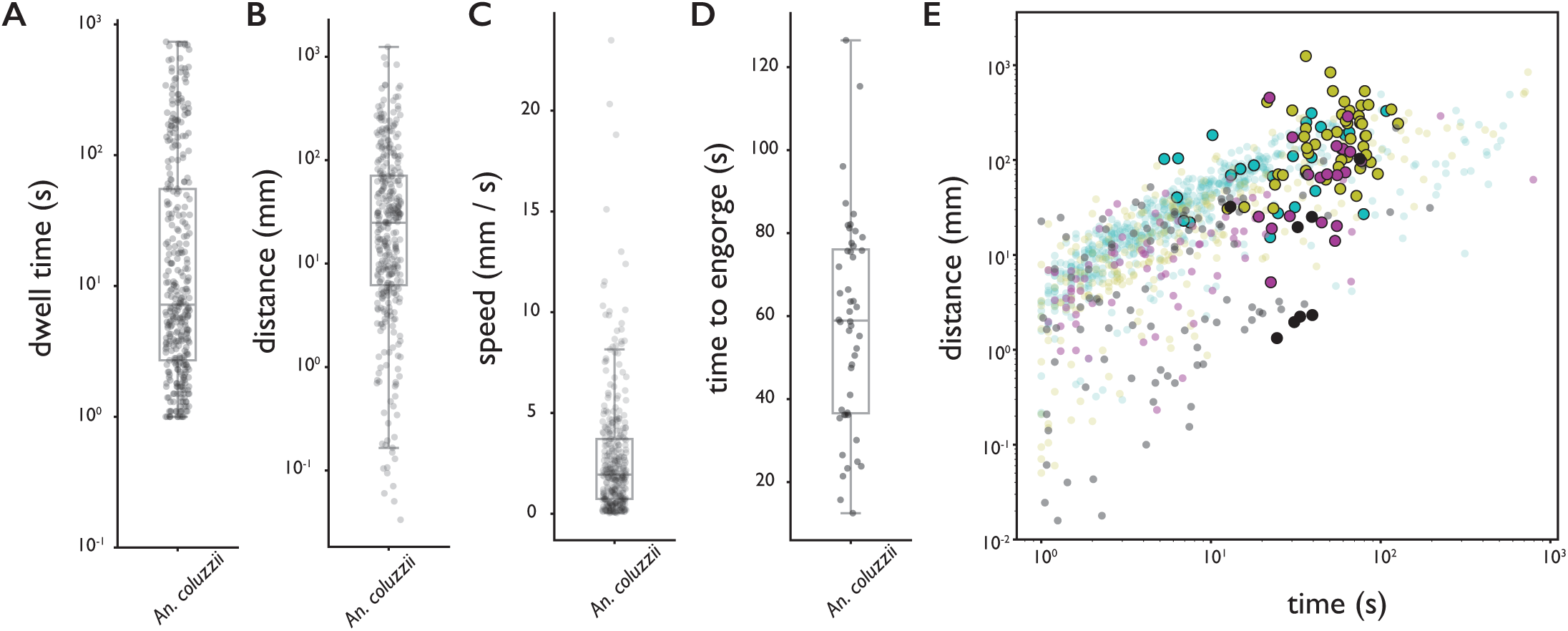
Behavioral statistics of *An. coluzzii* (A-D) and all four species (E). Each datapoint is derived from an individual trajectory, boxes indicate quartiles. (**A**) The time spent on the bite surface. (**B**) The total distance covered walking on the surface during a trajectory. (**C**) The mean velocity during a trajectory. (**D**) The time from landing to full engorgement (for trajectories leading to full engorgement). (**E**) The duration of a trajectory (total time for trajectories not leading to engorgement (transparent dots), time to full engorgement for trajectories that led to full engorgement (opaque circles)) versus the distance covered during that trajectory. The different colors denote different species, *Ae. aegypti*: magenta, *Ae. albopictus*: black, *An. stephensi*: cyan, *An. coluzzii*: yellow.

Fig. 2E shows the time spent on the surface versus the distance covered for trajectories that did (large opaque circles) and did not (small transparent dots) lead to full engorgement for the four species. As expected, rather short trajectories do not lead to engorgement, yet less intuitive is the observation that exploratory trajectories that do not lead to engorgement rarely exceed the duration of successful feeding trajectories (8% of non-feeding trajectories takes longer than the mean time to engorge). This suggests that a mosquito’s search for blood has a characteristic timescale that is independent of success, and when blood is not found within the time a typical meal takes, the search is aborted.

We further explored this observation using individual *Ae. albopictus* which were offered a bite substrate with a meal of PBS with or without ATP. As PBS alone does not lead to engorgement, mosquitoes offered the PBS only feeder never engorged whereas mosquitoes interacting with the PBS + ATP feeder engorged to full repletion in the majority of cases (55%). High resolution trajectory analysis enables us to dissect behavioral patterns that lead to (non-)feeding; a trajectory here is defined as landing, the ensuing behavioral sequence, followed by leaving the bite substrate by walking or flying (see Videos 8 and 9 for two example trajectories). Figure 3 presents ethograms of *Ae. albopictus* on these two bite substrates, and in agreement with the data in Fig. 2E, shows that trajectories on feeders without ATP (non-feeding) have an approximately equal maximum duration as trajectories leading to full engorgement on the feeder with ATP. While mosquitoes do not increase the duration of exploratory trajectories when not feeding to repletion, the number of exploratory bouts mosquitoes undertook on the PBS only substrate was significantly higher compared to the PBS + ATP case (Wilcoxon rank-sum test *p* < 0.05), resulting in a slightly longer total exploration time (Fig. 3C). This suggests that mosquitoes not finding their desired resource increase the frequency with which they initiate searches rather than the duration of individual searches. This observation may be interpreted in the context of the dangers associated with blood-feeding: while on a host, a mosquito runs the risk of being noticed and subsequently killed. When not finding blood, it may therefore be beneficial to abort the search and evacuate from a risky, yet unproductive situation to try elsewhere. Figure 3 furthermore shows a strong behavioral heterogeneity between individual mosquitoes. While all individuals are from the same mosquito population (and raised and maintained under identical conditions) and interact with the same bite substrate, there is a clear heterogeneity in the number of times a mosquito visits the surface, the amount of time she spends exploring the surface, and the behaviors they engage in. Automatic classification of locomotion behaviors, shows that some individuals often land on the surface to engage in short interactions, while other individuals undertake much longer trajectories. These long trajectories, in turn, vary in the amount of stationary versus locomotion behaviors. The richness of these data highlight the potential of the biteOscope to quantitatively characterize the intricacy of individual behaviors hidden in population averages.

**Figure 3.**
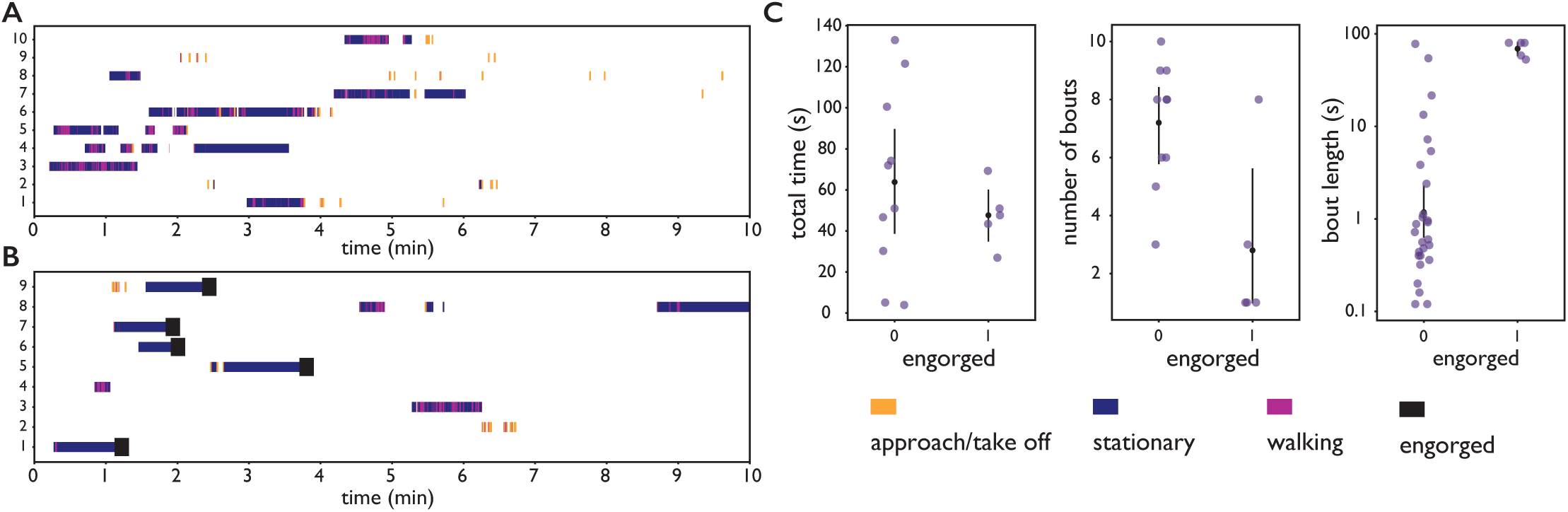
Feeding behavior of individual *Ae. albopictus*. (**A, B**) Ethograms of individual *Ae. albopictus* interacting with a bite substrate offering a PBS only meal (A) and a meal consisting of PBS + 1 mM ATP (B). Distinct exploratory bouts appear as continuous blocks in the ethogram and are labelled according to the behavior being displayed: flight (yellow), walking (purple), and stationary (dark blue), engorgement to full repletion is marked by a black box. (**C**) Behavioral statistics of the data displayed in A and B showing the total time spent on the bite substrate (left), the number of exploratory bouts undertaken (middle), and the length of individual bouts (right), of *Ae. albopictus* exploring the PBS only substrate (labelled 0) and those that engorged to full repletion on the PBS + ATP substrate (labelled 1). Individual data points are shown in purple, the mean and associated 95% confidence interval are depicted by a black dot and bar, respectively. Individuals that were offered the PBS + ATP substrate but did not feed to full repletion were excluded from this analysis.

### Pose estimation, behavioral classification, and contact-dependent sensing

We next turned to body part tracking to acquire a more detailed description of behavioral trajectories. Body part tracking is powerful to address a variety of questions, e.g. by determining points of surface contact of specific appendages, or to estimate the pose of an animal, which when tracked over time can be translated into a behavioral sequence. We used a recently developed deep learning framework, DeepLabCut (***Mathis et al., 2018***), to train a convolutional neural network (CNN) to detect the head, proboscis, abdomen, abdominal tip, and six legs of *Ae. aegypti* and *Ae. albopictus*. Due to their morphological similarity, the same CNN can be used to track the body parts of both *Aedes* species with a mean accuracy of 10 pixels (250 micrometer) in a 4.3 × 4.3 cm field of view.

Figure 4A-C shows body part tracking results of *Ae. albopictus* and reveals the choreography of three distinct behaviors. Anterior grooming is characterized by circular motion of the forelegs followed by the proboscis, while the middle legs remain stationary (see Video 10). During walking, the tips of all six legs oscillate along the body axis while the proboscis explores laterally (see Video 11), while during probing, the fore and middle legs pull towards the body and the proboscis remains stationary (see Video 12). Inference is done on raw images and the obtained coordinates thus subject to movement of the mosquito. To correct for this, the coordinates are translated and rotated to align along the body axis taking the abdominal tip as the origin. Figure 4D-I shows time series of the obtained egocentric coordinates and their corresponding wavelet transforms. The three behaviors each are associated with distinct periodic movements: smooth periodic motion of the forelegs during anterior grooming (x, and y coordinates), punctuated oscillations along the body axis during walking (x coordinate), and faster jerky movement during probing (x, and y coordinate of forelegs, y coordinate of middle legs). These trajectories can be used in concert with locomotion and body-shape features as inputs for behavioral classification algorithms. The data outputted by our computational pipeline is ideally suited for classification in either a supervised (e.g. (***Kain et al., 2013***; ***Kabra et al., 2013***)) or unsupervised (e.g. (***Berman et al., 2014***; ***Pereira et al., 2019***)) approaches.

**Figure 4.**
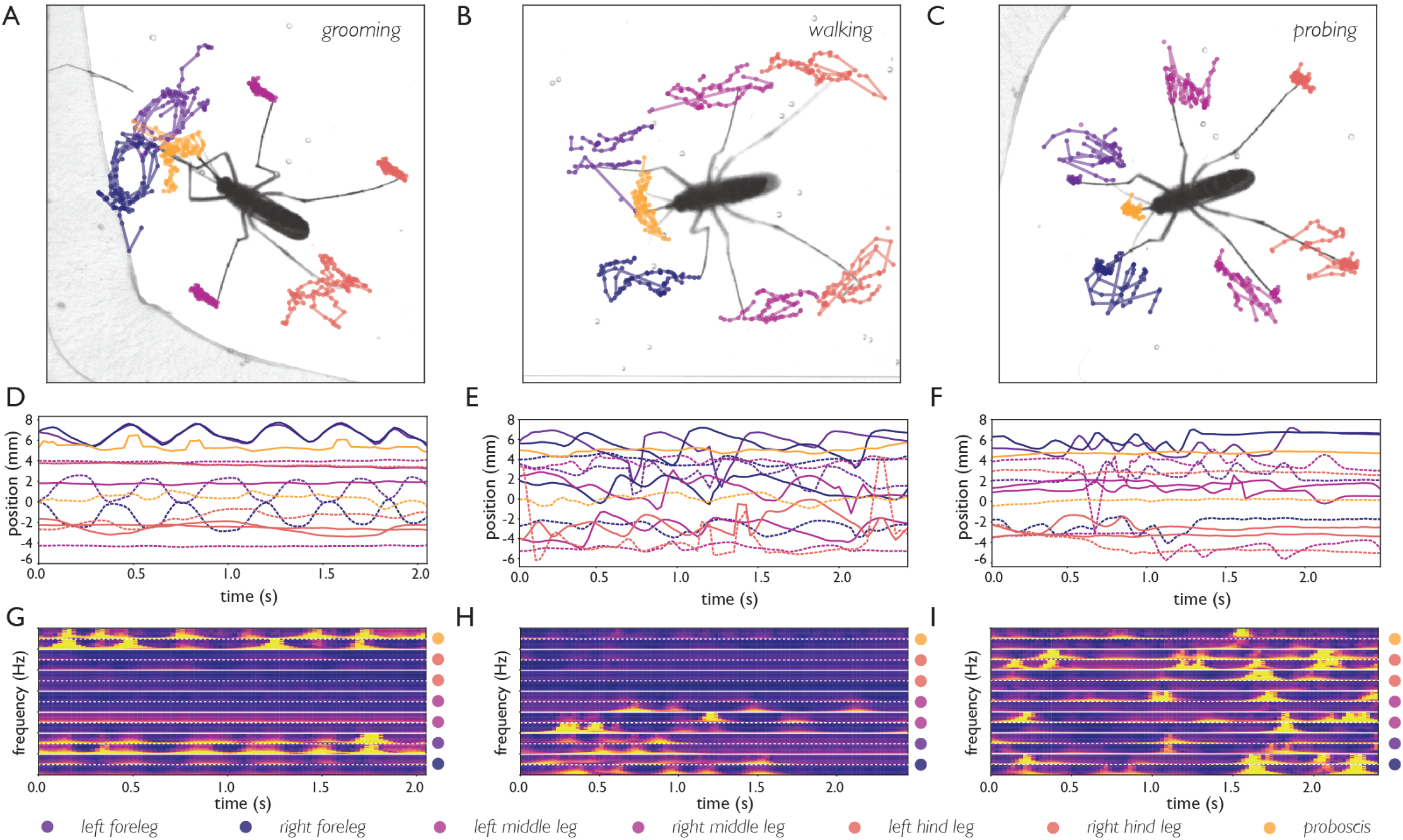
Body part tracking reveals movement patterns of specific behaviors. Color coding of plots in panels A-F are displayed at the bottom of the figure. (**A - C**) Trajectories of the tips of the six legs and proboscis of an *Ae. albopictus* female grooming her antennae (A), walking (B), and probing (C). (**D-F**) Time traces showing egocentric x (full lines) and y (dashed lines) coordinates of the body parts of mosquitoes shown in A-C. Anterior grooming is characterized by smooth periodic movement in the x and y planes. During walking the x-coordinate shows a swing that alternates between fore, middle, and hind leg; probing shows rapid pulling of the fore and middle legs towards the body. (**G-I**) Continuous wavelet transforms of the body part coordinates highlight the periodicity of movements. The amplitude of the spectrogram is indicated by the color, going from low (purple) to high (yellow). Yellow bands indicate periodic movement of a body part. Spectrograms of the 7 body parts are stacked and separated by white lines (color coding on the right shows stacking order, with the x-coordinate of the body part on top, and y-coordinate on the bottom (x, and y coordinates are separated by a dashed line)).

#### DEET repels *An. coluzzii* upon contact with legs

Next, we explored the use of body part tracking within the context of contact-dependent sensing by *An. coluzzii. Anopheles* and *Aedes* mosquitoes have an overall similar body plan, yet the length of their maxillary palps (an olfactory appendage projecting from the head) is very different with anophelines having maxillary palps with a length comparable to the proboscis, while *Aedes* palps are much shorter. We therefore trained a CNN for *Anopheles* body parts, which additionally tracks the position of the maxillary palps (mean accuracy for *Anopheles* body parts: 8 pixels, 200 micrometer).

Through this approach, we addressed a recently posed hypothesis stating that *An. coluzzii* may be repelled upon contact with N,N-diethyl-meta-toluamide (DEET) (***Afify et al., 2019***). Although DEET has been in use as an effective insect repellent for decades, its mode of action remains poorly understood. Afify *et al* reported that *An. coluzzii* is not capable of smelling DEET (i.e. detecting volatile DEET through its antennae), yet DEET may prevent *An. coluzzii* from locating humans by masking odorants emanating from potential hosts. However, it remained an open question if contact of the tarsal sensilla on *An. coluzzii’s* legs is a second mode through which DEET repels *An. coluzzii*.

We addressed this question by imaging *An. coluzzii* offered a bite substrate partly coated with DEET. Figure 5 shows that *An. coluzzii* do land on both the DEET coated and uncoated surface and there is a moderate decrease in landing rate on DEET coated portion (the landing rate is 1.9 times lower). The time *An. coluzzii* spend on the DEET coated surface, however, is much shorter: trajectories on the DEET coated surface (*n* = 34) are on average 7 times shorter when compared to the uncoated surface (*n* = 412). Furthermore, the longest residence time observed on the DEET coated surface was less than 6 seconds, whereas individual *An. coluzzii* spent up to 52 seconds on the uncoated surface. From these data we conclude that *An. coluzzii* do approach and land on the DEET coated surface, but avoid (prolonged) contact with it, indicating that *An. coluzzii* indeed is not strongly repelled by volatile DEET at very close range, yet avoids it on contact.

**Figure 5.**
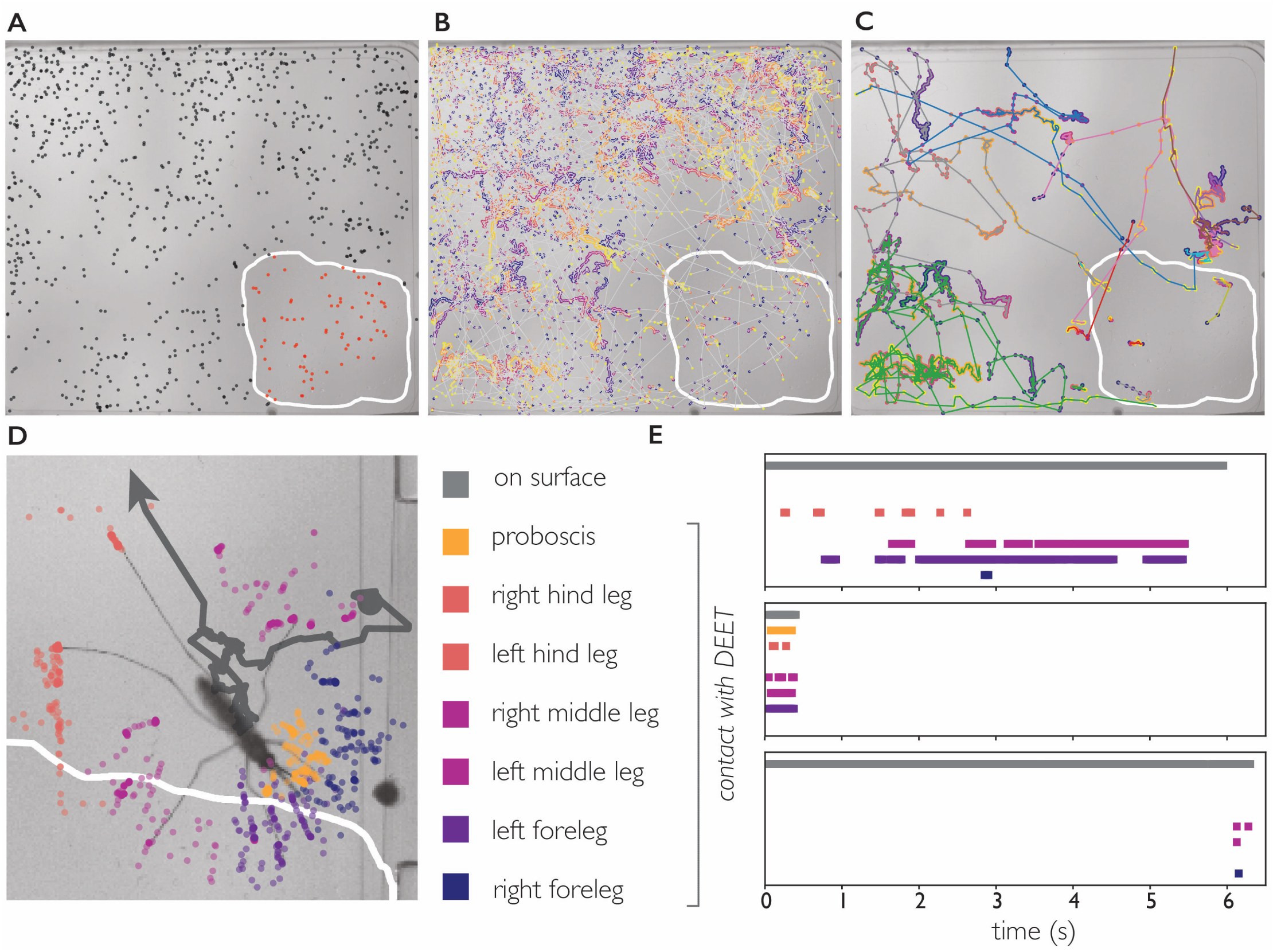
DEET repels *An. coluzzii* on contact with legs. (**A**) Landings on a substrate partly coated with DEET (white line indicates DEET coated surface). Black dots indicate landings outside the DEET area, red dots indicate landings inside the DEET area. The landing rate in the DEET area is approximately 1.9 times lower compared to the non-treated surface. (**B**) Trajectories of mosquito movement on the surface. Dots of individual tracks are colored from purple (start of the track) to yellow (end of the track). *An. coluzzii* on average spend 7 times longer on the non-coated surface compared to the DEET coated surface. (**C**) Example tracks of mosquitoes landing on the non-treated area and subsequently entering the DEET coated area. (**D**) Body part tracking of a mosquito near the edge of the DEET coated surface. The grey line indicates the movement of the center of mass of the mosquito (a dot indicates the start of the track, arrowhead departure). Colored dots indicate the position of the legs and proboscis during the section of the trajectory where the mosquito is within reach of the DEET coated area (indicated by the white line). (**E**) Ethogram showing typical behavioral patterns when a mosquito comes in contact with DEET. The grey bar (top) indicates that a mosquito is anywhere on the surface (including the uncoated area), the colored bars indicate contact of a specific appendage with DEET. The top panel corresponds to the mosquito shown in (D) illustrating a mosquito that walks towards the DEET area, contacts it with several legs, and flies away. The middle panel is an example of ‘touch and go’ contact in which a mosquito lands on the DEET area, contacts it with several legs and proboscis, and takes off. The bottom panel shows a mosquito that after a long exploratory bout outside the DEET area, takes off as soon as the right foreleg and both middle legs contact the DEET area.

We next asked what appendages mediate this contact dependent avoidance. The 34 trajectories in which *An. coluzzii* visited the DEET area consisted of 25 ‘touch and go’ events in which an individual approached the DEET surface in flight, landed, and immediately took off after first contact (residence time on DEET surface < 0.5 second, see Video 13 for a typical ‘touch and go’ event played at 1/4 speed). In the remaining 9 trajectories, *An. coluzzii* landed outside the DEET area and moved onto it (see Fig. 5 and video 14). We performed body part tracking on these trajectories and developed analysis software that scores how often a specific body part visits an arbitrarily shaped region of interest. We observed that the legs of individuals came in contact with the DEET surface in all cases, whereas the proboscis only came in contact with the DEET surface in 5/9 cases (in the non-contact cases the entire proboscis remained outside the boarders of the DEET treated area). Together, these observations demonstrate that *An. coluzzii* are indeed repelled upon contact with DEET, and indicate that this behavior is mediated by sensilla on the legs, and likely not the proboscis. This is in agreement with observations in *Ae. aegypti* which are also repelled by DEET upon leg contact and not by proboscis contact (***Dennis et al., 2019***). However, in contrast to *An. coluzzii, Ae. aegypti* can also detect the volatile form of DEET through its antennae (***DeGennaro et al., 2013***). While DEET is a potent repellent for a broad range of insects, it is interesting to see that certain modes of action are conserved across species, while others are not. The observation that both *Ae. aegypti* and *An. coluzzii* avoid DEET upon leg contact, while only the former can detect the volatile form, may guide efforts aimed at uncovering the underlying molecular mechanism.

## Discussion

The biteOscope provides an alternative for current methods using human subjects or mice to study mosquito blood feeding. The elimination of the need for a human subject opens new avenues of research, e.g. allowing blood feeding studies with pathogen infected mosquitoes, enabling precise surface manipulations and characterization of the associated behavior, and facilitates the use of high-resolution imaging and machine learning based image analysis. Through these innovations, the biteOscope increases experimental throughput and expands the type of experiments that can be performed and measurements that can be made.

We used the biteOscope to describe behavioral patterns of 4 medically relevant mosquito species and anticipate that such datasets will provide a useful ‘behavioral baseline’ for future studies quantifying the effect of a mosquito’s physiology on blood feeding behavior. The role of pathogen infections is particularly interesting in this respect, as several lines of evidence suggest that infections may alter feeding behavior, e.g. by affecting the structural integrity of the salivary glands or other body parts, or inducing systemic change through the immune system or infection of neural tissues (***Rossignol et al., 1984***; ***Girard et al., 2007***; ***Cator et al., 2013***; ***Turley et al., 2009***). A quantitative understanding of such behavioral alterations, however, is lacking. Gaining such insights is of high epidemiological relevance, as mathematical models suggest that (pathogen induced) changes in bite behavior can have important implications for pathogen transmission (***Cator et al., 2014***; ***Abboubakar et al., 2016***). In addition to pathogen induced behavioral changes, there are many other promising lines of inquiry, including the behavioral influence of the microbiome (***Dickson et al., 2017***), and assessment of the behavioral effects of population intervention strategies aimed at curbing pathogen transmission, such as *Wolbachia* infection or mosquitoes that are genetically engineered to be refractory to infection. The biteOscope is well positioned to address such questions.

We developed computational tools that allow the behavioral monitoring of mosquitoes at an unprecedented level of detail. Body part tracking of *Ae. albopictus* revealed specific movement patterns underlying distinct behaviors, and by tracking the individual body parts of *An. coluzzii*, we discovered that they are repelled by DEET upon leg contact—a mechanism that may work in concert with other ways in which DEET prevents anopheline mosquitoes to locate humans. Our results suggest a minor role for the proboscis in mediating DEET avoidance, and highlight the use of body part tracking in assigning roles to the various sensory appendages the mosquito body has. The recent surge in genetic tools available to manipulate mosquitoes is shedding light on the genetic elements that mediate pathogen transmission relevant behaviors (***Matthews et al., 2019***; ***Ingham et al., 2019***; ***Greppi et al., 2020***). Combining such molecular level insights with detailed behavioral tracking and chemical surface patterning, may enable a deep understanding of how contact dependent sensing drives blood feeding, and other important phenotypes such as insecticide resistance and egg laying preferences.

The biteOscope is designed with a variety of possible users in mind. It has a relatively modest price tag (900 - 3500 USD depending on the configuration), uses readily available materials and components, and when disassembled fits in a backpack—characteristics we hope will facilitate adoption. Beyond the lab, we foresee interesting applications of the behavioral tracking of mosquitoes in (semi-)field settings, and expect that innovative tools that provide high-quality quantitative data will enable discoveries in this space. We anticipate that the techniques and computational tools presented here will provide a fresh perspective on mosquito behaviors that are relevant to pathogen transmission, and enable researchers to gain a detailed understanding of blood feeding without having to sacrifice their own skin.

## Methods and Materials

### Mosquito rearing and maintenance

The mosquito species/strains used in this study are described in Supplementary Table S1. Larvae were hatched and reared in water at a density of approximately 200 larvae per liter on a diet of fish food. Adult mosquitoes were maintained at 28 °C, 75% relative humidity, and a photoperiod of 12 hours light : 12 hours dark in 30 cm^3^ screened cages having continuous access to 10% sucrose. Prior to experiments, mosquitoes were deprived of sucrose for 6-12 hours while having access to water. Mosquitoes aged 6 - 25 days old were used for behavioral experiments. Experiments using *Ae. aegypti* and *Ae. albopictus* were performed during light hours, while experiments with *An. stephensi* and *An. coluzzii* were performed during dark hours. Mosquitoes had no access to water during experiments.

### biteOscope hardware

A full list of components necessary to build the biteOscope is available in Supplementary Table S2. Depending on the experimental requirements, several components can be easily adapted (e.g. cage geometry or bite substrate) or replaced by more economical alternatives (e.g. imaging components).

#### Cage, bite substrate, and environmental control

Cages were constructed from 1/16 inch thick clear cast acrylic sheets (McMaster Carr) cut to the required dimensions using a laser cutter (Epilog). To facilitate mounting of the bite substrate, an opening having the same dimensions as the bite substrate was cut in the floor or one of the walls of the cage (all design files are available on Github). We noted that orientation of the bite substrate affects both the landing rate of mosquitoes (e.g. *Ae. albopictus* had a lower landing rate on vertically mounted substrates compared to those mounted in the floor) and their orientation (on vertical surfaces mosquitoes aligned with gravity, head up bottom down). While this suggests that orientation is an interesting parameter to explore, all experiments presented here were performed with floor-mounted substrates to prevent behavioral biases possibly associated with vertically mounted substrates. The bite substrate was made using a 70 mL culture flask (Falcon 353109) filled with warm water maintained at 37 °C by a Raspberry Pi taking the input of a waterproof temperature probe (DS18b20, Adafruit) to control a Peltier element (digikey) used for heating. If desired, the same Raspberry Pi can operate a 12 Volt solenoid valve (Adafruit) to control the inflow of gas. An artificial meal of phosphate buffered saline (sigma-aldrich) (supplemented with 1 mM of adenosine triphosphate (sigma aldrich) where noted) was applied to the rectangular section of the outside of the culture flask and covered with a Parafilm membrane. This creates a fluid cell supported by the membrane and the outside of the culture flask. The artificial meal is maintained at 37 °C by the water inside the flask.

#### Imaging and illumination

Images were acquired at 25 or 40 frames per second using a Basler acA2040-90um camera controlled using Pylon 5 software running on an Ubuntu 18.04 computer (NUC8i7BEH). The camera was equipped with a 100 mm macro lens (Canon macro EF 100mm f/2.8L). Illumination for *Aedes* experiments was provided by two white light LED arrays (Vidpro LED-312), while IR LEDs (Taobao) were used for *Anopheles* experiments. The same camera was used for white light and IR illuminated experiments. Thorlabs components were used to arrange all optical components and the experimental cage at suitable distance.

### Computational tools

All image processing and downstream analysis code was written in Python 3 and is available from Github (https://github.com/felixhol). Raw images were background subtracted, thresholded, and subjected to a series of morphological operations to yield binary images representing mosquito bodies of which the center of mass was determined using SciPy (***Virtanen et al., 2019***). The Crocker– Grier algorithm (***Crocker and Grier, 1996***) was used to link the obtained coordinates belonging to an individual mosquito in time using trackPy (***Allan et al., 2016***). The obtained tracking data is used to select all images that make up a single behavioral trajectory (e.g. landing, exploration, feeding, and take off) and store cropped image sequences centered on the focal mosquito. Cropped images are used to determine a mosquito’s body shape at each timepoint of a trajectory to subsequently infer engorgement status by computationally removing all appendages and fitting an active contour model (using OpenCV (***Bradski, 2000***)) to the remaining body shape.

The DeepLabCut framework (***Mathis et al., 2018***) was used to train a convolutional neural network (ResNet architecture) to detect the most distal part of the 6 legs, the abdominal tip, the center of the abdomen, the head, the tip of the proboscis, and for anophelines the tip of the maxillary palps. Due to their similar appearance, *Ae. aegypti* and *Ae. albopictus* can be analyzed using the same network, while a second network was trained for *An. stephensi* and *An. coluzzii*. Approximately 350 images were used to train the *Aedes* dataset, while approximately 300 images were used for the *Anopheles* dataset yielding an accuracy of 10 pixels (250 micrometer) and 8 pixels (200 micrometers) in a 4.3 × 4.3 cm field of view, respectively. Cropped image sequences (described above) were used for inference. To facilitate downstream analysis of body part tracking data, all body part coordinates were aligned along the body axis (defined along the abdominal tip and center of the abdomen) yielding coordinates invariant of body orientation or movement.

### Experiment specific procedures

#### Feeding experiments

Population experiments (Figures 1 and 2) were performed with 15 - 30 individuals in a 10 cm^3^ cage. Groups of mosquitoes were recorded for up to 1 hour and replaced by a new group for a subsequent recording. Individual *Ae. albopictus* females (Fig. 3) were recorded for 10 minutes per mosquito and discarded after the experiment. Movement status (Fig. 3 A and B) was classified using the velocity derived from tracking.

#### DEET experiments

As DEET dissolves Parafilm and plastics, a glass surface was placed on top of the heated cell culture flask (no artificial meal was present during DEET experiments). The glass surface was partly coated with 50% N,N-diethyl-meta-toluamide (DEET) using a cotton swab. Groups of 20 *An. coluzzii* females (14 days old) were released into a 10 cm^3^ cage with the DEET coated substrate mounted in the floor. Images were acquired at 40 frames per second for 1 hour. Mosquito and body part tracking was performed as described above.

## Acknowledgments

We thank Haripriya Vaidehi Narayanan, Shailabh Kumar, William Gilpin, Hongquan Li, Leslie Vosshall, Veronica Jove, and all members of the Prakash and Lambrechts labs for valuable discussions; and Catherine Lallemand, Sylvain Golba, and Patricia Baldacci for help with the rearing of mosquitoes. *An. stephensi* strain Sda500 and *An. coluzzii* strain N’Gousso were provided by the Centre de Production et Infection d’Anopheles of Institut Pasteur; *Ae. aegypti* strain Liverpool was provided by Leslie Vosshall (Rockefeller University); *Ae. aegypti* strain D2S3 (NR-45838) was provided by Centers for Disease Control and Prevention for distribution by BEI Resources, NIAID, NIH. FJHH was supported by a Rubicon fellowship for the Netherlands Foundation for Scientific Research, a Career Award at the Scientific Interface from the Burroughs Wellcome Fund, and a Marie Curie Fellowship from the European Union. L.L. is supported by the French Agence Nationale de la Recherche (grants ANR-16-CE35-0004-01 and ANR-18-CE35-0003-01), and the French Government’s Investissement d’Avenir program Laboratoire d’Excellence Integrative Biology of Emerging Infectious Diseases (grant ANR-10-LABX-62-IBEID). MP was supported by NIH DP2-AI124336 New Innovator Award and USAID Grand Challenges: Zika and Future Threats Award.

## Author contributions

Felix JH Hol, Conceptualization, Methodology, Software, Data curation, Formal analysis, Funding acquisition, Validation, Investigation, Visualization, Project administration, Writing—original draft, Writing—review and editing; Louis Lambrechts, Resources, Project administration, Writing—review and editing; Manu Prakash, Conceptualization, Funding acquisition, Project administration, Writing— review and editing

## Supplementary Material

**Video 1.** *Ae. albopictus* female landing, probing, and feeding to full repletion. Upon landing, the mosquito walks/explores the substrate for a short period to pierce the surface and insert her stylets, clearly visible as a flexible needle. The video shows a fast pulling motion of the fore and hind legs towards the body which is typical during the probing phase. While engorging, the body remains nearly motionless and the abdomen dilates visibly.

**Video 2.** An *Ae. aegypti* female lands, probes (visible as a pulling motion towards the body), walks several millimeters, probes again, and finally starts to engorge. Engorgement is clearly visible as a dilation of the abdomen. Video playing in real time.

**Video 3.** Several *An. stephensi* females explore the bite substrate, two feed to repletion. The individual that initiates feeding in the top right corner of the frame stops engorging half way, and subsequently moves to the left side of the frame to continue engorging. Video playing in real time.

**Video 4.** Several *An. coluzzii* explore interact with the bite substrate, two feed to repletion. Both *Anopheles* species quickly concentrate the obtained meal by excreting liquid (visible as a growing excretion droplet), *Aedes* excrete small droplets as well, yet to a much smaller extent. Video playing in real time.

**Video 5.** The stylet of an *Ae. aegypti* female evaluates the artificial meal it finds after piercing the membrane. The stylet is a flexible organ that bends, extends, and retracts in the liquid. Video playing in real time.

**Video 6.** Tracking the centroid of *Ae. aegypti*. The color of the centroid and the trail is a measure for the instantaneous velocity of the animal.

**Video 7.** The abdomen of an *Ae. aegypti* female expands dramatically during blood feeding. Fitting an active contour model to the mosquito body (after computationally removing appendages) provides the abdomen width (and other shape parameters) which can be used to estimate engorgement status.

**Video 8.** *Ae. albopictus* female walking onto the bite substrate (artificial meal of PBS without ATP), probing the substrate several times, and moving away. Video playing in real time.

**Video 9.** *Ae. albopictus* female exploring the surface of PBS only feeder (without ATP). While walking, the proboscis often moves laterally and taps the surface. Video playing in real time.

**Video 10.** Body part tracking of *Ae. albopictus* anterior grooming, corresponding to Fig. 4A, D, G. Video playing in real time. Grooming of the antennae and proboscis is characterized by a circular motion of the forelegs, and corresponding movement of the head. The middle and hind legs remain stationary during this behavior.

**Video 11.** Body part tracking of *Ae. albopictus* walking, corresponding to Fig. 4B, E, H. Video playing in real time. The stride sequence of *Ae. albopictus*, in which fore, middle, and hind legs swing in sequence, is visible.

**Video 12.** Body part tracking of *Ae. albopictus* probing, corresponding to Fig. 4C, F, I. Video playing in real time. While probing, all 4 species show rapid pulling motion of the legs towards the head (here a typical video of *Ae. albopictus* is shown). After the membrane is pierced, the location of the proboscis remains stationary.

**Video 13.** *An. coluzzii* landing on the DEET coated surface and immediately taking off. Video playing 4 times slower than real time. The majority of trajectories in which *An. coluzzii* comes into contact with the DEET coated surface results in an immediate take off.

**Video 14.** *An. coluzzii* moving onto the DEET coated surface. Body part tracking shows that this female lands outside the DEET coated area and subsequently her left fore and middle leg come into contact with the DEET coated portion. After a short contact, the mosquito flies away.

**Figure S1.**
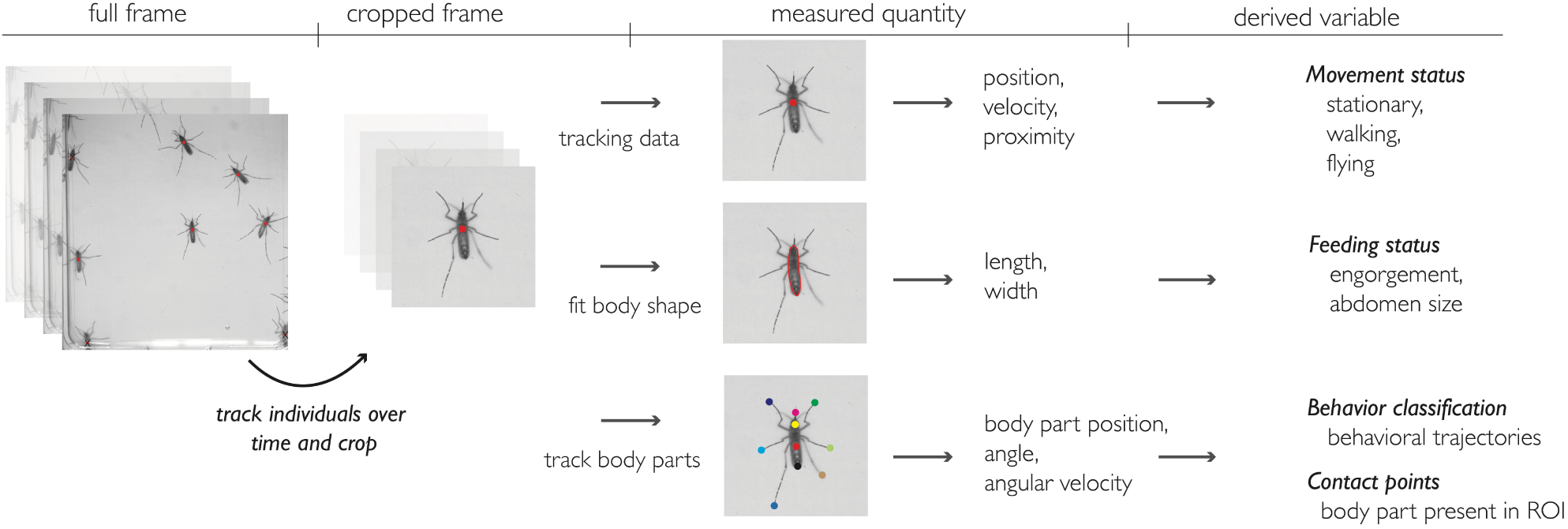
Overview of the computational pipeline.

**Figure S2.**
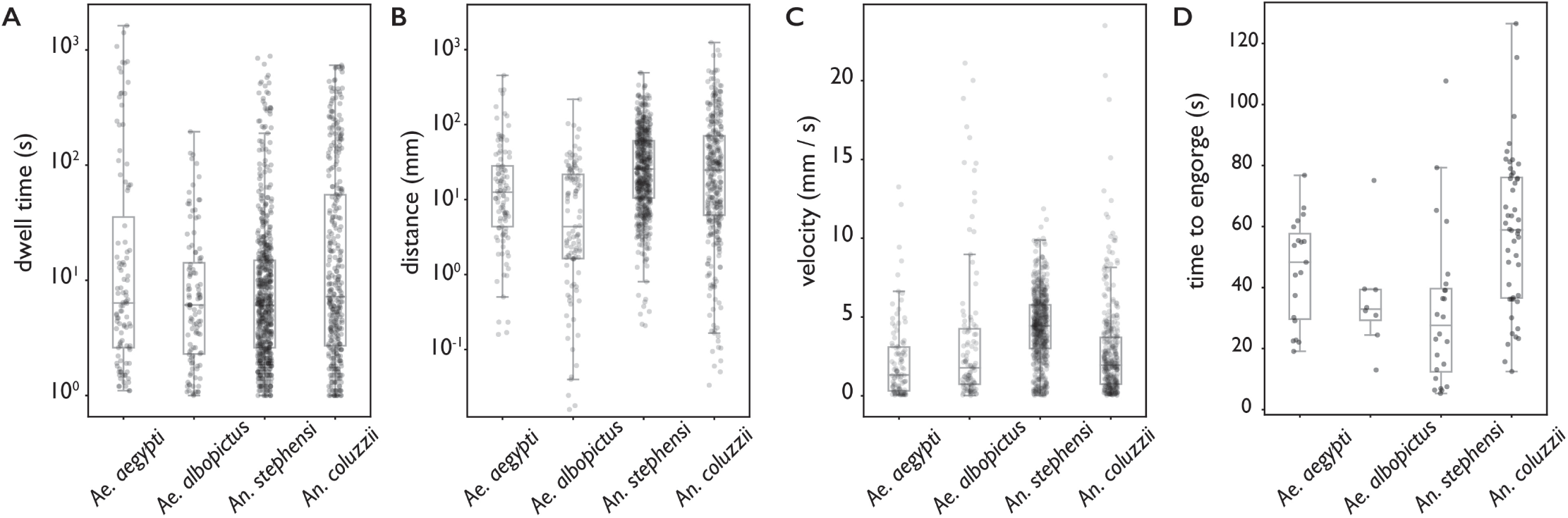
Behavioral statistics of *Ae. aegypti, Ae. albopictus, An. stephensi*, and *An. coluzzii*. Each datapoint is derived from an individual trajectory. (**A**) The time spent on the bite surface. (**B**) The total distance covered walking on the surface during this time. (**C**) The mean velocity during a trajectory. (**D**) The time from landing to full engorgement (for trajectories leading to full engorgement).

**Table S1.** Mosquito colonies used in this study.

| Species | Strain | Geographic origin | Generation | Source |
| --- | --- | --- | --- | --- |
| *Ae. aegypti* | KPPTN | Thailand | F 18 | Lambrechts lab, Institut Pasteur |
| *Ae. aegypti* | D2S3 | Puerto Rico x Nigeria cross | laboratory strain | BEI resources |
| *Ae. aegypti* | Liverpool | West Africa | laboratory strain | Vosshall lab, Rockefeller University |
| *Ae. albopictus* | BP | Vietnam | F 23 | Lambrechts lab, Institut Pasteur |
| *An. stephensi* | Sda500 | Pakistan | laboratory strain | CEPIA, Institut Pasteur, Paris |
| *An. coluzzii* | N'Gousso | Cameroon | laboratory strain | CEPIA, Institut Pasteur, Paris |

**Table S2.**
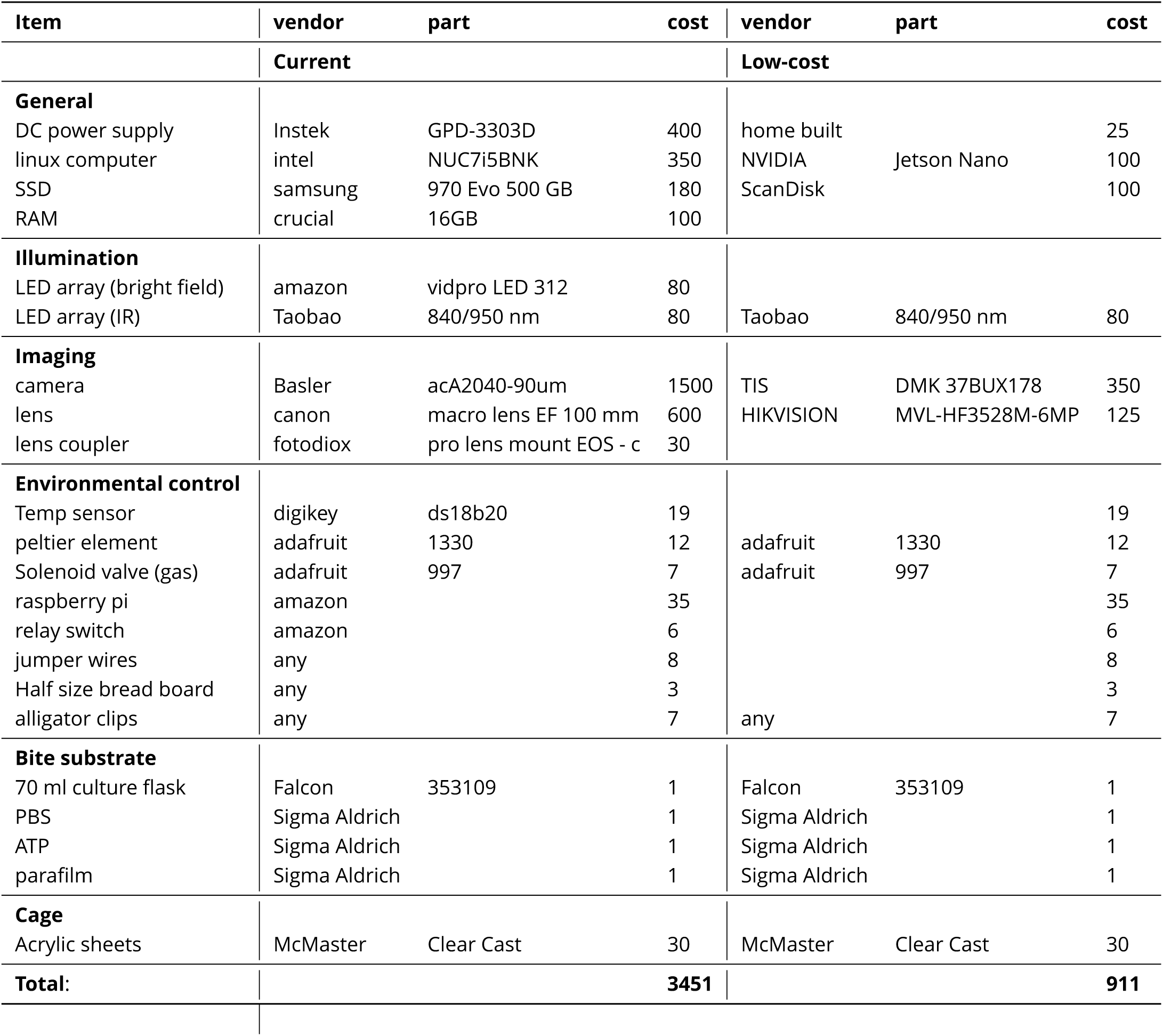
BiteOscope parts list. The left columns describe the set-up as used for all experiments described in the paper, the right columns describe a low-cost alternative. Vendors are suggestions, costs are in US dollars and approximate.

## References

Abboubakar H, Buonomo B, Chitnis N. Modelling the effects of malaria infection on mosquito biting behaviour and attractiveness of humans. Ricerche di matematica. 2016; 65(1):329–346.

Afify A, Betz JF, Riabinina O, Lahondère C, Potter CJ. Commonly used insect repellents hide human odors from Anopheles mosquitoes. Current Biology. 2019;.

Allan D, Caswell T, Keim N, van der Wel C. trackpy: Trackpy v0. 3.2. Zenodo http://doi.org/105281/zenodo. 2016; 60550:2019.

Benton R. The neurobiology of gustation in insect disease vectors: progress and potential. Current opinion in insect science. 2017; 20:19–27.

Berman GJ, Choi DM, Bialek W, Shaevitz JW. Mapping the stereotyped behaviour of freely moving fruit flies. Journal of The Royal Society Interface. 2014; 11(99):20140672.

Bradski G. The opencv library. Dr Dobb’s J Software Tools. 2000; 25:120–125.

Branson K, Robie AA, Bender J, Perona P, Dickinson MH. High-throughput ethomics in large groups of Drosophila. Nature methods. 2009; 6(6):451.

van Breugel F, Riffell J, Fairhall A, Dickinson MH. Mosquitoes use vision to associate odor plumes with thermal targets. Current Biology. 2015; 25(16):2123–2129.

Cator LJ, George J, Blanford S, Murdock CC, Baker TC, Read AF, Thomas MB. ‘Manipulation’without the parasite: altered feeding behaviour of mosquitoes is not dependent on infection with malaria parasites. Proceedings of the Royal Society B: Biological Sciences. 2013; 280(1763):20130711.

Cator LJ, Lynch PA, Thomas MB, Read AF. Alterations in mosquito behaviour by malaria parasites: potential impact on force of infection. Malaria journal. 2014; 13(1):164.

Choumet V, Attout T, Chartier L, Khun H, Sautereau J, Robbe-Vincent A, Brey P, Huerre M, Bain O. Visualizing non infectious and infectious Anopheles gambiae blood feedings in naive and saliva-immunized mice. PLoS One. 2012; 7(12):e50464.

Clements AN. The physiology of mosquitoes: International series of monographs on pure and applied biology: Zoology, vol. 17. Elsevier; 2013.

Corfas RA, Vosshall LB. The cation channel TRPA1 tunes mosquito thermotaxis to host temperatures. Elife. 2015; 4:e11750.

Crocker JC, Grier DG. Methods of digital video microscopy for colloidal studies. Journal of colloid and interface science. 1996; 179(1):298–310.

De Jong R, Knols B. Selection of biting sites on man by two malaria mosquito species. Experientia. 1995; 51(1):80–84.

DeGennaro M, McBride CS, Seeholzer L, Nakagawa T, Dennis EJ, Goldman C, Jasinskiene N, James AA, Vosshall LB. orco mutant mosquitoes lose strong preference for humans and are not repelled by volatile DEET. Nature. 2013; 498(7455):487.

Dekker T, Cardé RT. Moment-to-moment flight manoeuvres of the female yellow fever mosquito (Aedes aegypti L.) in response to plumes of carbon dioxide and human skin odour. Journal of Experimental Biology. 2011; 214(20):3480–3494.

Dennis EJ, Goldman OV, Vosshall LB. Aedes aegypti Mosquitoes Use Their Legs to Sense DEET on Contact. Current Biology. 2019; 29(9):1551–1556.

Dickson LB, Jiolle D, Minard G, Moltini-Conclois I, Volant S, Ghozlane A, Bouchier C, Ayala D, Paupy C, Moro CV, et al. Carryover effects of larval exposure to different environmental bacteria drive adult trait variation in a mosquito vector. Science advances. 2017; 3(8):e1700585.

Duvall LB, Ramos-Espiritu L, Barsoum KE, Glickman JF, Vosshall LB. Small-molecule agonists of Ae. aegypti neuropeptide Y receptor block mosquito biting. Cell. 2019; 176(4):687–701.

Galun R, Avi-Dor Y, Bar-Zeev M. Feeding response in Aedes aegypti: stimulation by adenosine triphosphate. Science. 1963; 142(3600):1674–1675.

Geier M, Boeckh J. A new Y-tube olfactometer for mosquitoes to measure the attractiveness of host odours. Entomologia experimentalis et applicata. 1999; 92(1):9–19.

Girard YA, Schneider BS, McGee CE, Wen J, Han VC, Popov V, Mason PW, Higgs S. Salivary gland morphology and virus transmission during long-term cytopathologic West Nile virus infection in Culex mosquitoes. The American journal of tropical medicine and hygiene. 2007; 76(1):118–128.

Gordon R, Lumsden W. A study of the behaviour of the mouth-parts of mosquitoes when taking up blood from living tissue; together with some observations on the ingestion of microfilariae. Annals of Tropical Medicine & Parasitology. 1939; 33(3-4):259–278.

Greppi C, Laursen WJ, Budelli G, Chang EC, Daniels AM, van Giesen L, Smidler AL, Catteruccia F, Garrity PA. Mosquito heat seeking is driven by an ancestral cooling receptor. Science. 2020; 367(6478):681–684.

Griffiths R, Gordon R, et al. An apparatus which enables the process of feeding by mosquitoes to be observed in the tissues of a live rodent; together with an account of the ejection of saliva and its significance in Malaria. Annals of tropical medicine and parasitology. 1952; 46(4):311–19.

Hagan RW, Didion EM, Rosselot AE, Holmes CJ, Siler SC, Rosendale A J, Hendershot JM, Elliot KS, Jennings EC, Nine GA, et al. Dehydration prompts increased activity and blood feeding by mosquitoes. Scientific reports. 2018; 8(1):6804.

Healy T, Copland M, Cork A, Przyborowska A, Halket J. Landing responses of Anopheles gambiae elicited by oxocarboxylic acids. Medical and Veterinary Entomology. 2002; 16(2):126–132.

Ingham VA, Anthousi A, Douris V, Harding NJ, Lycett G, Morris M, Vontas J, Ranson H. A sensory appendage protein protects malaria vectors from pyrethroids. Nature. 2019; p. 1–5.

Itskov PM, Moreira JM, Vinnik E, Lopes G, Safarik S, Dickinson MH, Ribeiro C. Automated monitoring and quantitative analysis of feeding behaviour in Drosophila. Nature communications. 2014; 5:4560.

Jones JC, Pilitt DR. Blood-feeding behavior of adult Aedes aegypti mosquitoes. The Biological Bulletin. 1973; 145(1):127–139.

Jové V, Gong Z, Hol FJH, Zhao Z, Sorrells TR, Carroll TS, Prakash M, McBride CS, Vosshall LB. The taste of blood in mosquitoes. bioRxiv. 2020; (954206).

Kabra M, Robie AA, Rivera-Alba M, Branson S, Branson K. JAABA: interactive machine learning for automatic annotation of animal behavior. Nature methods. 2013; 10(1):64.

Kain J, Stokes C, Gaudry Q, Song X, Foley J, Wilson R, de Bivort B. Leg-tracking and automated behavioural classification in Drosophila. Nature communications. 2013; 4(1910).

Lee R. Structure and function of the fascicular stylets, and the la bral and cibarial sense organs of male and female Aedes aeg ypti (L.)(Diptera, Culicidae). Quaestiones entomologicae. 1974;.

Mathis A, Mamidanna P, Cury KM, Abe T, Murthy VN, Mathis MW, Bethge M. DeepLabCut: markerless pose estimation of user-defined body parts with deep learning. Nature neuroscience. 2018; 21(9):1281.

Matthews BJ, Younger MA, Vosshall LB. The ion channel ppk301 controls freshwater egg-laying in the mosquito Aedes aegypti. eLife. 2019; 8:e43963.

McMeniman CJ, Corfas RA, Matthews BJ, Ritchie SA, Vosshall LB. Multimodal integration of carbon dioxide and other sensory cues drives mosquito attraction to humans. Cell. 2014; 156(5):1060–1071.

Moreira JM, Itskov PM, Goldschmidt D, Baltazar C, Steck K, Tastekin I, Walker SJ, Ribeiro C. optoPAD, a closed-loop optogenetics system to study the circuit basis of feeding behaviors. eLife. 2019; 8:e43924.

Moreira LA, Saig E, Turley AP, Ribeiro JM, O’Neill SL, McGraw EA. Human probing behavior of Aedes aegypti when infected with a life-shortening strain of Wolbachia. PLoS neglected tropical diseases. 2009; 3(12):e568.

Pereira TD, Aldarondo DE, Willmore L, Kislin M, Wang SSH, Murthy M, Shaevitz JW. Fast animal pose estimation using deep neural networks. Nature methods. 2019; 16(1):117–125.

Ribeiro J. Blood-feeding in mosquitoes: probing time and salivary gland anti-haemostatic activities in representatives of three genera (Aedes, Anopheles, Culex). Medical and veterinary entomology. 2000; 14(2):142–148.

Rossignol P, Ribeiro J, Spielman A. Increased intradermal probing time in sporozoite-infected mosquitoes. The American journal of tropical medicine and hygiene. 1984; 33(1):17–20.

Sparks JT, Vinyard BT, Dickens JC. Gustatory receptor expression in the labella and tarsi of Aedes aegypti. Insect biochemistry and molecular biology. 2013; 43(12):1161–1171.

Stanczyk NM, Mescher MC, De Moraes CM. Effects of malaria infection on mosquito olfaction and behavior: extrapolating data to the field. Current opinion in insect science. 2017; 20:7–12.

Turley AP, Moreira LA, O’Neill SL, McGraw EA. Wolbachia infection reduces blood-feeding success in the dengue fever mosquito, Aedes aegypti. PLoS Neglected Tropical Diseases. 2009; 3(9).

Vantaux A, de Sales Hien DF, Yaméogo B, Dabiré KR, Thomas F, Cohuet A, Lefèvre T. Host-seeking behaviors of mosquitoes experimentally infected with sympatric field isolates of the human malaria parasite Plasmodium falciparum: no evidence for host manipulation. Frontiers in Ecology and Evolution. 2015; 3:86.

Verhulst NO, Qiu YT, Beijleveld H, Maliepaard C, Knights D, Schulz S, Berg-Lyons D, Lauber CL, Verduijn W, Haasnoot GW, et al. Composition of human skin microbiota affects attractiveness to malaria mosquitoes. PloS one. 2011; 6(12):e28991.

Virtanen P, Gommers R, Oliphant TE, Haberland M, Reddy T, Cournapeau D, Burovski E, Peterson P, Weckesser W, Bright J, et al. SciPy 1.0–fundamental algorithms for scientific computing in Python. arXiv preprint 190710121. 2019;.

Werner-Reiss U, Galun R, Crnjar R, Liscia A. Sensitivity of the mosquito Aedes aegypti (Culicidae) labral apical chemoreceptors to phagostimulants. Journal of insect physiology. 1999; 45(7):629–636.

Zermoglio PF, Robuchon E, Leonardi MS, Chandre F, Lazzari CR. What does heat tell a mosquito? Characterization of the orientation behaviour of Aedes aegypti towards heat sources. Journal of insect physiology. 2017; 100:9–14.

